# Differential effects of Wnt-β-catenin signaling in Purkinje cells and Bergmann glia in SCA1

**DOI:** 10.1101/2021.10.21.465333

**Authors:** Kimberly Luttik, Leon Tejwani, Hyoungseok Ju, Terri Driessen, Cleo Smeets, Janghoo Lim

**Author notes:** Corresponding author: Dr. Janghoo Lim, 295 Congress Avenue, BCMM 154E, New Haven CT 06510. **Author Contributions:** K.L., L.T., H.J., and J.L. conceived and designed the study. K.L., L.T., H.J., T.D., and C.S. performed the experiments. K.L., L.T., H.J., T.D., and C.S. analyzed the data. K.L., L.T., and J.L. wrote the manuscript. All authors reviewed the manuscript and provided comments. **Competing interest statement:** The authors declare no competing interest.

## Abstract

Spinocerebellar ataxia type 1 (SCA1) is a dominantly inherited neurodegenerative disease characterized by progressive ataxia and degeneration of specific neuronal populations, including Purkinje cells (PCs) in the cerebellum. Previous studies have demonstrated a critical role for various evolutionarily conserved signaling pathways in cerebellar patterning, such as the Wnt-β-catenin pathway; however, the roles of these pathways in adult cerebellar function and cerebellar neurodegeneration are largely unknown. In this study, we found that Wnt-β-catenin activity was progressively enhanced in multiple cell types in the adult SCA1 mouse cerebellum, and that activation of this signaling occurs in an ataxin-1 polyglutamine (polyQ) expansion-dependent manner. Genetic manipulation of the Wnt-β-catenin signaling pathway in specific cerebellar cell populations revealed that activation of Wnt-β-catenin signaling in PCs alone was not sufficient to induce SCA1-like phenotypes, while its activation in astrocytes including Bergmann glia (BG) resulted in gliosis and disrupted BG localization, which was replicated in SCA1 mouse models. Our studies identify a novel mechanism in which polyQ-expanded ataxin-1 positively regulates Wnt-β-catenin signaling, and demonstrate that different cell types have distinct responses to the enhanced Wnt-β-catenin signaling in the SCA1 cerebellum, underscoring an important role of BG in SCA1 pathogenesis.

**Significance statement:** The mechanisms underlying the degeneration of specific cellular populations in various neurodegenerative disorders remain unknown. Here, we show that the polyQ expansion of ataxin-1 activates the Wnt-β-catenin signaling pathway in various cell types, including Purkinje cells and Bergmann glia, in the cerebellum of SCA1 mouse models. We used conditional mouse genetics to activate and silence this pathway in different cell types and found elevated activity of this signaling pathway impacted Bergmann glia and Purkinje cell populations differently. This study highlights the important role of Wnt-β-catenin signaling pathway in glial cell types for SCA1 pathogenesis.

## Introduction

Spinocerebellar ataxia type 1 (SCA1) is an adult-onset neurodegenerative disorder caused by a trinucleotide repeat expansion of a glutamine-encoding CAG tract in *ATXN1*^1^ In SCA1, specific neuronal populations degenerate at later stages of disease, including cerebellar Purkinje cells (PCs), brainstem cranial nerve nuclei, and inferior olive neurons^2^. Although ataxia-related motor changes typically manifest during adulthood, animal models of SCA1 have revealed substantial molecular and circuit-level alterations in the cerebellum at time points prior to the onset of robust behavioral deficits^3–8^, suggesting developmental abnormalities can contribute to long-term cerebellar health and SCA1 pathogenesis. Furthermore, although *ATXN1* is ubiquitously expressed throughout the brain^9–11^, the cellular and molecular mechanisms leading to the selective degeneration of specific cell types is largely unknown.

Among the different signaling pathways that have been identified through unbiased profiling of ataxia animal models, several studies suggest that components of the Wnt signaling pathway are perturbed in SCA1 and other forms of ataxia, with ataxin-1 and ataxin-3 null mice exhibiting altered expression of genes involved in the Wnt signaling pathway^8, 12–14^. The Wnt signaling pathway is comprised of three highly conserved signal transduction pathways: the canonical Wnt-β-catenin pathway, and the noncanonical Wnt-planar cell polarity and Wnt-calcium pathways^15^. Within the cerebellum, the canonical Wnt-β-catenin pathway plays crucial roles in regulating the proliferation, migration, and differentiation of diverse cell types during cerebellar morphogenesis^16–23^. Previous studies have shown that aberrant activation of Wnt-β-catenin signaling in cerebellar granule precursor cells can inhibit their proliferation, prompting precocious differentiation during development^21^. Additionally, a number of genes encoding key components in the Wnt-β-catenin signaling pathway, including *Apc, Gsk3β, Ctnnb1* (encoding β-catenin), and *Lef-1*, as well as its target genes, including *Ccnd1* and *Myc*, are expressed in adult PCs^9, 24, 25^. Together, these studies indicate that Wnt-β-catenin signaling is active and is homeostatically regulated in the cerebellum during development and into adulthood. Thus, although it is clear that Wnt plays a fundamental role in establishing proper cerebellar cytoarchitecture, the precise physiological role of persistent Wnt signaling in different cell types of the adult cerebellum remains elusive. Furthermore, the nature of Wnt-β-catenin signaling perturbation and the functional implications of this dysregulation in the SCA1 cerebellum at various stages of disease are unclear.

In this study, we demonstrate that Wnt-β-catenin signaling is activated in multiple cerebellar cell types in an age-dependent manner *in vivo* in SCA1 through both cell autonomous and non-cell autonomous mechanisms. We identified a novel molecular mechanism through which ataxin-1 positively regulates Wnt-β-catenin signaling in a polyglutamine (polyQ)-dependent manner to cell autonomously enhance Wnt target gene expression in PCs. Interestingly, expression of polyQ-expanded ataxin-1 specifically in PCs also resulted in increased production of multiple secreted Wnt ligands and higher Wnt activity in other cell populations, including Bergmann glia (BG). To understand the impact of Wnt signaling in these different cerebellar cell types, we used conditional mouse genetics approaches to manipulate levels of Wnt-β-catenin signaling in a cell-type specific manner in SCA1 mouse models. Our data revealed differential effects of Wnt-β-catenin signaling in different cell types, with perturbations in Wnt signaling in BG having a greater impact on overall cerebellar health than in PCs. Taken together, these data describe the effect of altering Wnt signaling in different cell types in the adult cerebellum and support a role for BG in the progressive cerebellar dysfunction observed in SCA1.

## Results

### Wnt-β-catenin signaling is enhanced in the SCA1 cerebellum

Due to the well-defined role of Wnt-β-catenin signaling (Figure 1A) in cerebellar circuit formation^16–23^, and the involvement of developmental changes in long-term cerebellar health in SCA^14, 5^, we sought to interrogate if and how Wnt-β-catenin signaling is involved in SCA1 at various stages of disease progression. To this end, we first examined Wnt-β-catenin signaling in SCA1 knock-in mice that express *Atxn1* containing 154 CAG repeats under the control of its endogenous promoter (*Atxn1*^154Q/2Q^; SCA1 KI^26^). Gene expression of Wnt-β-catenin target genes *Ccnd1* and *c-Myc* was upregulated *in vivo* in the SCA1 KI cerebellum in an age-dependent manner, with elevated expression at 30 weeks but not at 6 weeks (Figure 1B).

**Figure 1.**
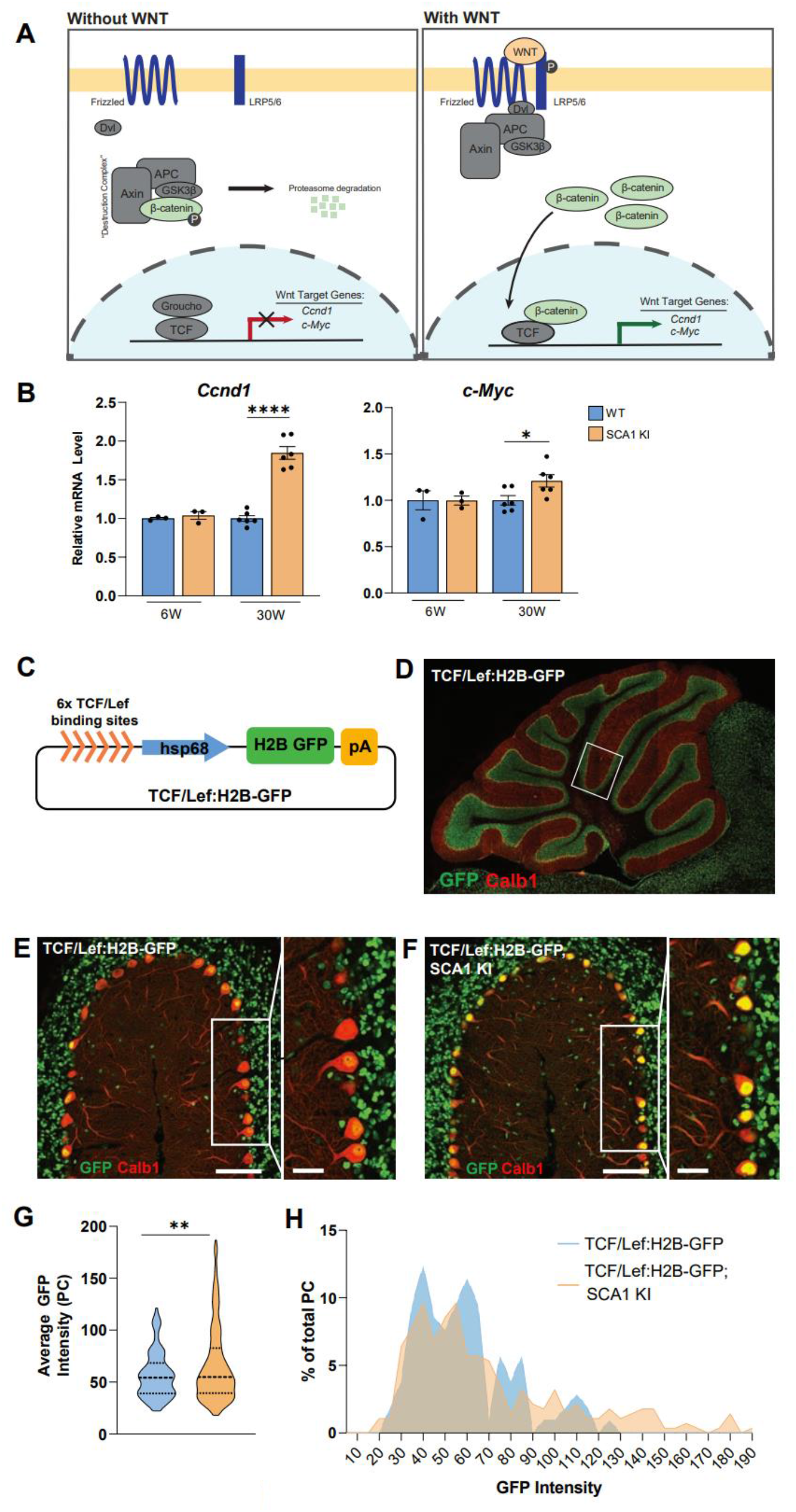
Enhanced activation of Wnt-β-catenin signaling in the SCA1 KI mouse cerebellum. **A,** Overview of Wnt-β-catenin signaling pathway. In the absence of Wnt ligands, β-catenin is degraded by a degradation complex. In the presence of Wnt ligands, β-catenin accumulates and translocates to the nucleus where it binds with TCF family transcription factors to activate Wnt-responsive genes, including *Ccnd1*, and *c-Myc*. **B,** RT-qPCR of Wnt-β-catenin target gene expression in SCA1 KI mouse cerebellum at 6 and 30 weeks, normalized to WT littermate controls (6 weeks, n=3 animals per genotype; 30 weeks, n=6 per genotype). **C,** Schematics of Wnt-β-catenin signaling reporter (TCF/Lef:H2B-GFP). **D,** Representative image of Wnt-β-catenin signaling reporter (TCF/Lef:H2B-GFP) cerebellum, stained with GFP (Wnt-β-catenin signaling activity) and Calbindin (Calb1, PCs). **E,F,** Representative images of 19 week TCF/Lef:H2B-GFP **(E)** and TCF/Lef:H2B-GFP; SCA1 KI **(F)** mouse cerebellar lobule 5, stained with GFP and Calb1, scale bar 100 μm, inset 25 μm. **G,H,** Quantification of intensity of Wnt-β-catenin signaling activity in PCs, as average GFP intensity in all Calb1^+^ cells **(G),** and as percentage of total PCs counted binned by GFP intensity **(H)** (TCF/Lef:H2B-GFP; SCA1 KI, n=2; TCF/Lef:H2B-GFP, n=1). \**P*<0.05, \*\**P*<0.01, \*\*\*\**P*<0.0001, by student’s *t*-test.

To determine the specific cell types in which Wnt signaling is activated in SCA1, we utilized a transgenic reporter mouse (TCF/Lef:H2B-GFP) for the Wnt-β-catenin signaling pathway that uses a minimal promoter containing six TCF binding sites to express a H2B-GFP fusion protein upon canonical Wnt activation (Figure 1C,D)^27^. We crossed SCA1 KI mice with TCF/Lef:H2B-GFP mice and examined reporter activity in the cerebellum at 19 weeks of age, a timepoint in which substantial gene expression changes have been reported (Figure 1E,F)^28^. Wild-type (WT) reporter animals showed a baseline level of Wnt signaling reporter activity in multiple cell types in adult cerebellum, including PCs (as indicated by the PC marker calbindin, calb1), as well as surrounding cell types in the granule cell layer and molecular layer (Figure 1D,E), supporting previous studies demonstrating that Wnt signaling persists into adulthood. SCA1 KI reporter mice exhibited an increased average GFP intensity (Figure 1F,G) and increased proportion of PCs with higher intensity of Wnt-β-catenin signaling reporter activity in the cerebellum (Figure 1F,H). Additionally, increase in reporter fluorescence was observed in other surrounding cell types in the molecular layer, PC layer, and granular cell layer of the SCA1 KI cerebellum relative to WT reporter animals (Figure 1E,F). Collectively, these results demonstrate that expression of polyQ-expanded ataxin-1 in the cerebellum increases canonical Wnt signaling activity in multiple different cerebellar cell types during SCA1 disease progression.

### Ataxin-1 positively regulates Wnt-β-catenin signaling in a polyQ-dependent manner

To determine the mechanism through which polyQ-expanded ataxin-1 affects the transcriptional output of Wnt-β-catenin signaling (Figure 2A), we utilized an established β-catenin-responsive luciferase reporter, superTOPFlash^29^ (Figure 2B), which is activated by diverse upstream activators, including Wnt ligands, Dishevelled, LiCl (inhibitor of GSK3β, a negative regulator of Wnt-β-catenin signaling), and β-catenin (Figure 2A). Transfection of polyQ-expanded ataxin-1 strongly enhanced superTOPFlash activity in HeLa cells treated with Wnt3A conditioned media (Figure 2C), confirming our *in vivo* finding that polyQ-expanded ataxin-1 is able to enhance Wnt-β-catenin signaling activity. To identify at which level(s) ataxin-1 modulates the Wnt-β-catenin signaling pathway, we performed similar luciferase reporter assays using Wnt-β-catenin signaling activators at different stages of the Wnt-β-catenin signaling cascade, including Dishevelled-3 transfection, LiCl treatment, and β-catenin transfection (Figure 2D-F). We observed that in the absence of active Wnt signaling, ataxin-1 alone did not alter superTOPFlash activation. However, in all cases in which Wnt signaling was active, ataxin-1 could further enhance the level of superTOPFlash activation (Figure 2C-F), suggesting that polyQ-expanded ataxin-1 likely operates in parallel with or downstream of β-catenin to enhance transcription of Wnt-β-catenin target genes in SCA1.

**Figure 2.**
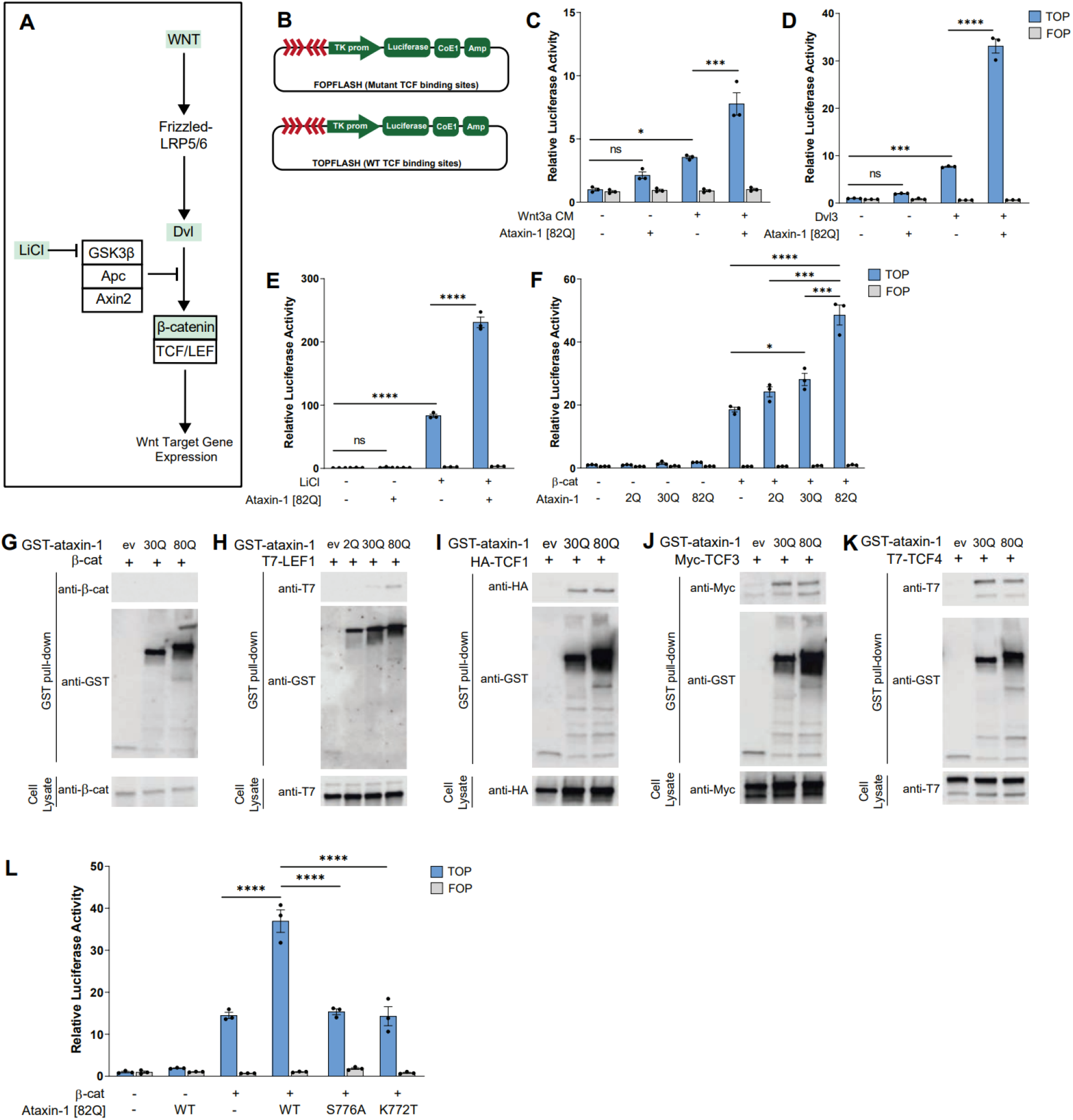
Ataxin-1 activates Wnt-β-catenin signaling transcription by binding to TCF family transcription factors. **A,** Wnt-β-catenin signaling cascade schematic, with components of the pathway targeted in Luciferase assay **(C-F,L)** highlighted in green. **B,** Schematic of TOPFlash, a β-catenin-responsive luciferase reporter, and FOPFlash, negative control, constructs, in which TCF binding sites trigger luciferase activity. **C-F**, Quantification of luciferase activity (TOPFlash) upon co-transfection of HeLa cells with ataxin-1 [82Q] with Wnt-β-catenin signaling activators, including **(C)** Wnt3A-conditioned media (Wnt3a CM), **(D)** Dishevelled 3 (Dvl3), **(E)** LiCl treatment, an inhibitor of GSK3β (negative regulator of Wnt signaling), and **(F)** β-catenin (β-cat) (n=3 per treatment group). **G-K,** Co-affinity purification assays of polyQ-expanded ataxin-1 of 30Q and 82Q length and TCF/β-catenin transcription factors in HeLa cells, including **(G)** β-catenin, **(H)** LEF1, **(I)** TCF1, **(J)** TCF3, **(K)** TCF4. Top panel shows expression of TCF/β-catenin transcription factors and GST-ataxin-1 after affinity purification on Glutathione-Sepharose 4B beads. Bottom panel shows total cell lysate. ev = GST-empty vector. **L,** PolyQ-expanded ataxin-1 enhances Wnt-β-catenin signaling while non-pathogenic forms of ataxin-1 do not in Luciferase assay. S776A is phosphorylation-defective, K772T is nuclear localization-defective (n=3 per treatment group). *P<0.05, ***P<0.001, ****P<0.0001, ns, non-significant, by one-way ANOVA with Tukey’s post-hoc analysis.

Transcriptional regulation of target genes in the canonical Wnt signaling pathway occurs via the coordination of β-catenin with several transcription factors following its translocation to the nucleus^30^. To further elucidate how ataxin-1 regulates Wnt-β-catenin signaling, we performed co-affinity purification experiments to determine whether ataxin-1 physically interacts with the various transcription factors involved in canonical Wnt signaling, including β-catenin and TCF/LEF family members. Although canonical Wnt signaling requires β-catenin as a co-activator of transcription, no physical interaction between ataxin-1 and β-catenin was observed (Figure 2G). However, a physical interaction between ataxin-1 and TCF/LEF family members LEF1 (HUGO name LEF1), TCF1 (HUGO name TCF7), TCF3 (HUGO name TCF7L1), and TCF4 (HUGO name TCF7L2) was observed (Figure 2H-K). Interestingly, the interaction between ataxin-1 and Lef1 increased in a polyQ-dependent manner (Figure 2H), providing a potential molecular mechanism for cell autonomous ataxin-1-mediated Wnt-β-catenin signaling activation in SCA1. Overall, these data demonstrate that mutant ataxin-1 can physically interact with multiple effectors of Wnt-β-catenin target gene transcription and, in the presence of β-catenin, increase expression of downstream genes.

The pathogenicity of ataxin-1 is dependent on several key features and domains of the protein, including polyQ expansion, serine 776-phosphorylation, and its nuclear localization signal^31, 32^. Therefore, we next investigated the involvement of these various features on Wnt induction by ataxin-1. First, longer ataxin-1 polyQ tract length resulted in increased superTOPFlash activation (Figure 2F). Additionally, polyQ-expanded ataxin-1 carrying mutations of a key phosphorylation site (S776A) or nuclear localization signal (K772T) abrogated the activation of Wnt-β-catenin signaling by ataxin-1 (Figure 2L). Because polyQ expansion, phosphorylation at S776, and the ability to localize to the nucleus are crucial for ataxin-1 pathogenicity, and modulating any of these properties mitigated the effect of ataxin-1 on Wnt-β-catenin signaling activation, these data reveal a novel mechanism through which pathogenic forms of ataxin-1 may contribute to transcriptional dysregulation in SCA1.

### PC-specific expression of polyQ-expanded ataxin-1 is sufficient to induce cerebellar Wnt-β-catenin hyperactivation

Next, to determine whether polyQ-expanded ataxin-1 can activate Wnt-β-catenin signaling through cell autonomous mechanisms in PCs *in vivo*, we utilized a PC-specific transgenic mouse model of SCA1 in which polyQ-expanded ataxin-1 with 63 glutamine repeats was overexpressed in PCs (SCA1 Tg [63Q]), which was originated due to a germline contraction event from SCA1 Tg [82Q] line^33^. Similar to SCA1 KI mice, transcript levels of Wnt target genes *Ccnd1* and *c-Myc* were elevated in the cerebellum of SCA1 Tg [63Q] mice in an age-dependent manner compared to WT controls (Figure 3A). Interestingly, protein levels of active β-catenin, but not mRNA levels of *Ctnnb1*, were also elevated in the cerebellum of 30-week SCA1 Tg [63Q] mice (Figure 3B-D), suggesting post-transcriptional regulation of β-catenin levels by ataxin-1. Furthermore, we measured significant increases in GFP intensity, corresponding to Wnt-β-catenin signaling reporter activity, in PCs of 12-week SCA1 Tg [63Q] mice crossed with TCF/Lef:H2B-GFP reporter mice, compared to control mice (Figure 3E-H). Surprisingly, we found that the increased activity of Wnt-β-catenin signaling reporter was not only limited cell autonomously within the PCs of SCA1 Tg [63Q] cerebellum, but also non-cell autonomously in other surrounding cell types in the PC layer, as well as molecular and granule cell layers (Figure 3E-F). Finally, we found that expression of certain secreted Wnt ligands was elevated at 30 weeks in SCA1 Tg [82Q] mice (Figure S1), which could contribute to the observed non-cell autonomous activation of Wnt signaling in surrounding cerebellar cell types. These data demonstrate that overexpression of polyQ-expanded ataxin-1 exclusively in PCs leads to enhanced activation of Wnt-β-catenin signaling cell autonomously in PCs, as well as surrounding cell types of the SCA1 cerebellum through indirect, non-cell autonomous mechanisms.

**Figure 3.**
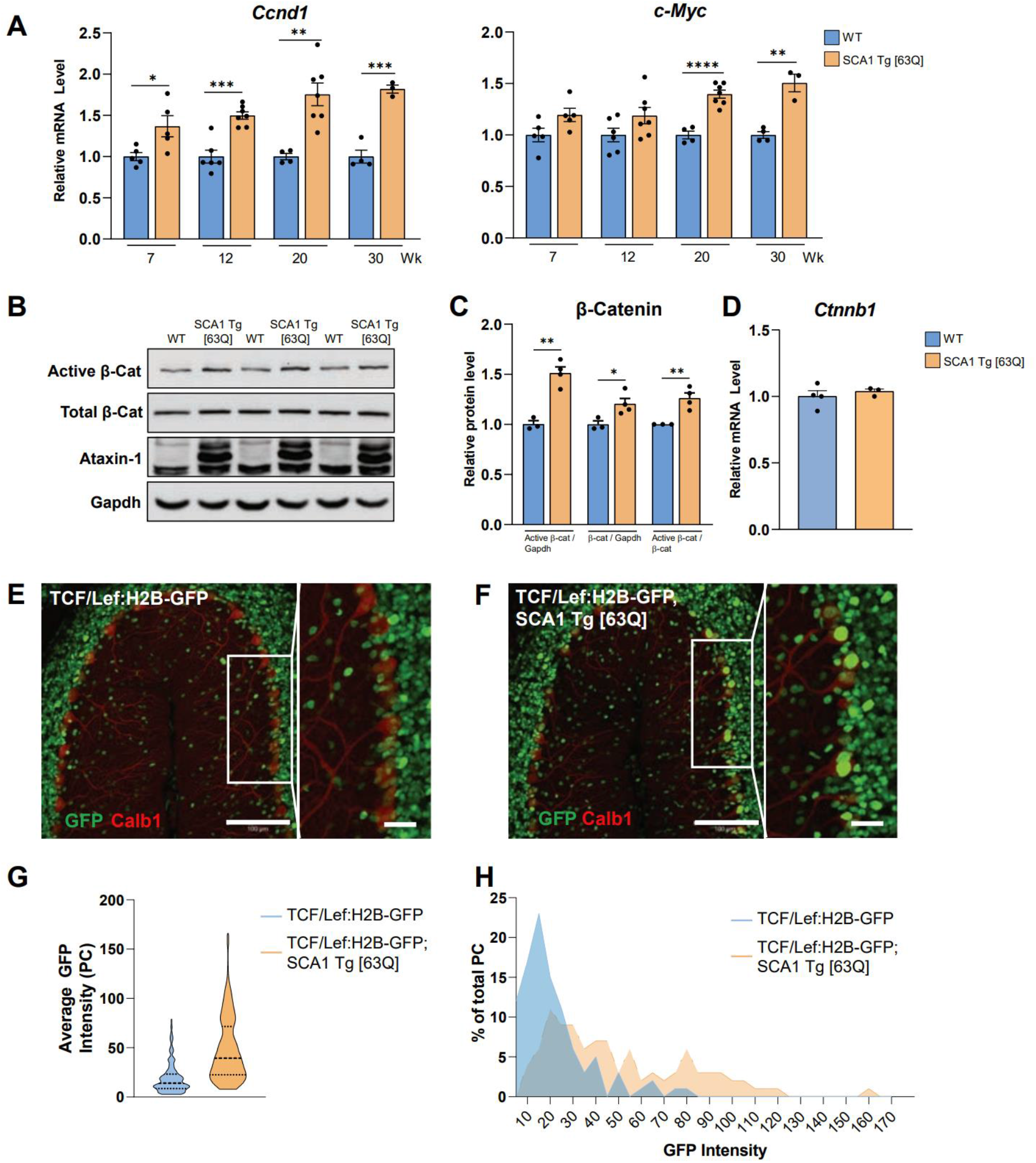
Enhanced activation of Wnt-β-catenin signaling in the SCA1 Tg [63Q] mouse cerebellum. **A,** RT-qPCR of Wnt-β-catenin target gene expression in SCA1 Tg [63Q] mouse cerebellum at 7, 12, 20, and 30 weeks, normalized to WT littermate controls (n=3-7 animals per genotype). **B,** Western blot images of β-catenin protein expression (total and active forms) and ataxin-1 protein expression in whole cerebellar extracts of WT and SCA1 Tg [63Q] mice at 30 weeks. Gapdh was used as a loading control. **C,** Quantification of total and active β-catenin protein expression levels from **(B)**, normalized to Gapdh and WT expression levels (WT, n=3; SCA1 Tg [63Q], n=4). **D,** RT-qPCR of *Ctnnb1* mRNA expression in SCA1 Tg [63Q] mouse cerebellum at 30 weeks, normalized to WT littermate controls (n=3 per genotype). **E,F,** Representative images of 12-week TCF/Lef:H2B-GFP **(E)** and SCA1 Tg [63Q]; TCF/Lef:H2B-GFP **(F)** mouse cerebellar lobule 5, stained with Calb1 and GFP. Scale bar 100 μm, inset 25 μm. **G,H**, Quantification of intensity of Wnt-β-catenin signaling activity in PCs, as average GFP intensity in all Calb1^+^ cells **(G),** and as percentage of total PCs counted binned by GFP intensity **(H)** (TCF/Lef:H2B-GFP; SCA1 Tg [63Q], n=4; TCF/Lef:H2B-GFP, n=3). *P<0.05, **P<0.01, ***P<0.001, ****P<0.0001, by student’s *t*-test.

### Genetic manipulation of canonical Wnt signaling in PCs has minimal impact on cerebellar health and PC survival

To determine whether enhanced Wnt-β-catenin signaling in PCs of SCA1 animals directly leads to neurodegeneration or is secondary to disease progression, we utilized multiple genetic approaches to conditionally suppress or activate Wnt-β-catenin signaling in a cell-type specific manner in WT and SCA1 mice (Figures 4, 5). We first assessed whether inhibition of Wnt-β-catenin signaling in PCs was able to rescue pathological deficits in SCA1 through conditional deletion of *Ctnnb1*, the gene encoding β-catenin, specifically in PCs (*Ctnnb1* PC cKO; *Ctnnb1^fl/fl^; Pcp2-cre* mice) of both WT and SCA1 Tg [63Q] animals (Figure 4A). Immunostaining confirmed successful removal of β-catenin in PCs of *Ctnnb1* PC cKO mice, with surrounding cells still maintaining β-catenin expression (Figure 4B). Decreased levels of β-catenin protein were also confirmed in whole cerebellar extracts of *Ctnnb1* PC cKO mice (Figure 4C). Silencing Wnt-β-catenin signaling on a WT background did not significantly impact PC health during adulthood (Figure 4, Figure S2). As expected, molecular layer thickness was reduced in SCA1 Tg [63Q] mice; however, this was not rescued by inhibition of Wnt-β-catenin signaling in *Ctnnb1* PC cKO; SCA1 Tg [63Q] mice at 12, 20, or 30 weeks compared to littermate controls (Figure 4D,E). Additionally, there were no significant changes in climbing fiber innervation between *Ctnnb1* PC cKO; SCA1 Tg [63Q] and SCA1 Tg [63Q] mice at 12 or 20 weeks (Figure 4F,G). To confirm these results were not due to the reduced polyQ length or background, we also analyzed molecular layer thickness and climbing fiber innervation in an independent cohort of SCA1 transgenic animals overexpressing polyQ-expanded ataxin-1 with 82 repeats in PCs (SCA1 Tg [82Q])^33^. Similar to SCA1 Tg [63Q] mice, conditional deletion of *Ctnnb1* in PCs of SCA1 Tg [82Q] animals did not rescue molecular layer thickness or climbing fiber innervation deficits in mice analyzed at 21 weeks of age (Figure S2). Furthermore, silencing of Wnt-β-catenin signaling in PCs was not sufficient to decrease astrogliosis and microgliosis in SCA1 Tg [63Q] mice at 20 weeks, as shown by Gfap and Iba1 staining of astrocytes and microglia, respectively (Figure 4H-K). These data demonstrate that inhibition of Wnt-β-catenin signaling in PCs specifically was not sufficient to prevent pathological deficits observed in transgenic SCA1 mice.

**Figure 4.**
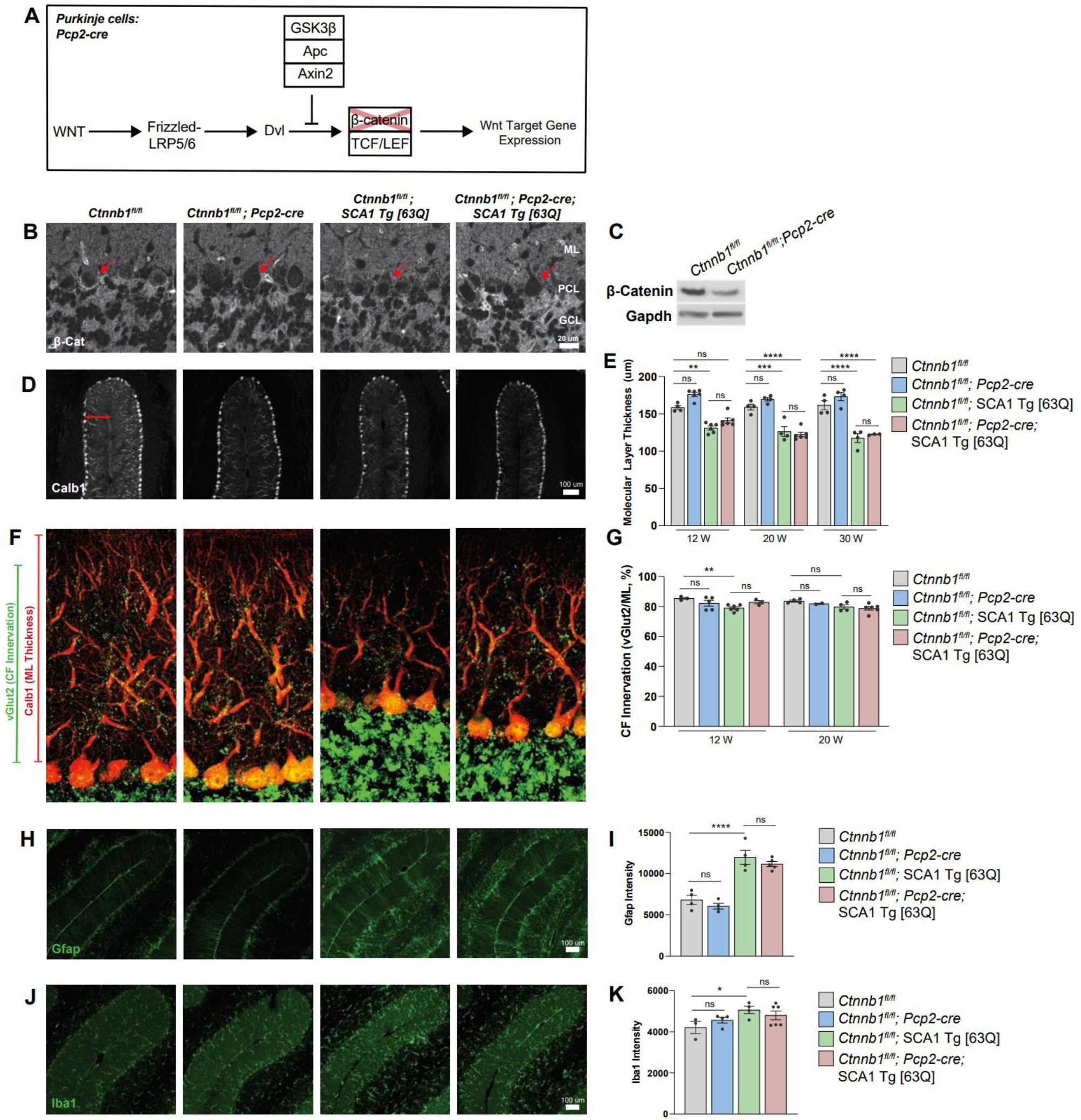
Loss of Wnt-β-catenin signaling in PCs does not prevent SCA1 phenotypes in SCA1 Tg [63Q] mice. **A,** Schematic of Wnt-β-catenin signaling silencing in PCs by *Ctnnb1* conditional deletion. **B,** Immunohistochemistry showing loss of β-catenin in the PCs of *Ctnnb1* PC cKO (*Ctnnb1*^fl/fl^; *Pcp2*-cre) mice, with arrows indicating PCs. **C,** Representative Western blot images of β-catenin protein expression in control and *Ctnnb1* PC cKO mice, with Gapdh as loading control. **D,E,** Representative images of Calb1 staining of 20 week cerebellar lobule 5 in control and *Ctnnb1* PC cKO mice on WT and SCA1 Tg [63Q] backgrounds **(D)**, to measure molecular layer thickness quantified at 12, 20, and 30 weeks in **(E)**, n=4, 6, 6, 6 (12 weeks), n=4, 4, 4, 6 (20 weeks), n=4, 4, 4, 3 (30 weeks)**. F,G,** Representative images of vGlut2 and Calb1 staining of 20-week cerebellar lobule 5 in control and *Ctnnb1* PC cKO mice on WT and SCA1 Tg [63Q] backgrounds **(F)**, to quantify climbing fiber (CF) innervation (ratio of vGlut2 / molecular layer thickness) at 12 and 20 weeks **(G),** n= 4, 6, 6, 6 (12 weeks), n= 4, 4, 4, 6 (20 weeks). **H,** Representative images of Gfap staining at 20 weeks in control and *Ctnnb1* PC cKO mice, on WT and SCA1 Tg [63Q] backgrounds, quantified in **(I),** n=4, 4, 4, 5. **J,** Representative images of Iba1 staining at 20 weeks in control and *Ctnnb1* PC cKO mice, on WT and SCA1 Tg [63Q] backgrounds, quantified in **(K),** n=3, 4, 4, 6. *P<0.05, **P<0.01, ***P<0.001, ****P<0.0001, ns, non-significant, by one-way ANOVA with Tukey’s post-hoc analysis.

**Figure 5.**
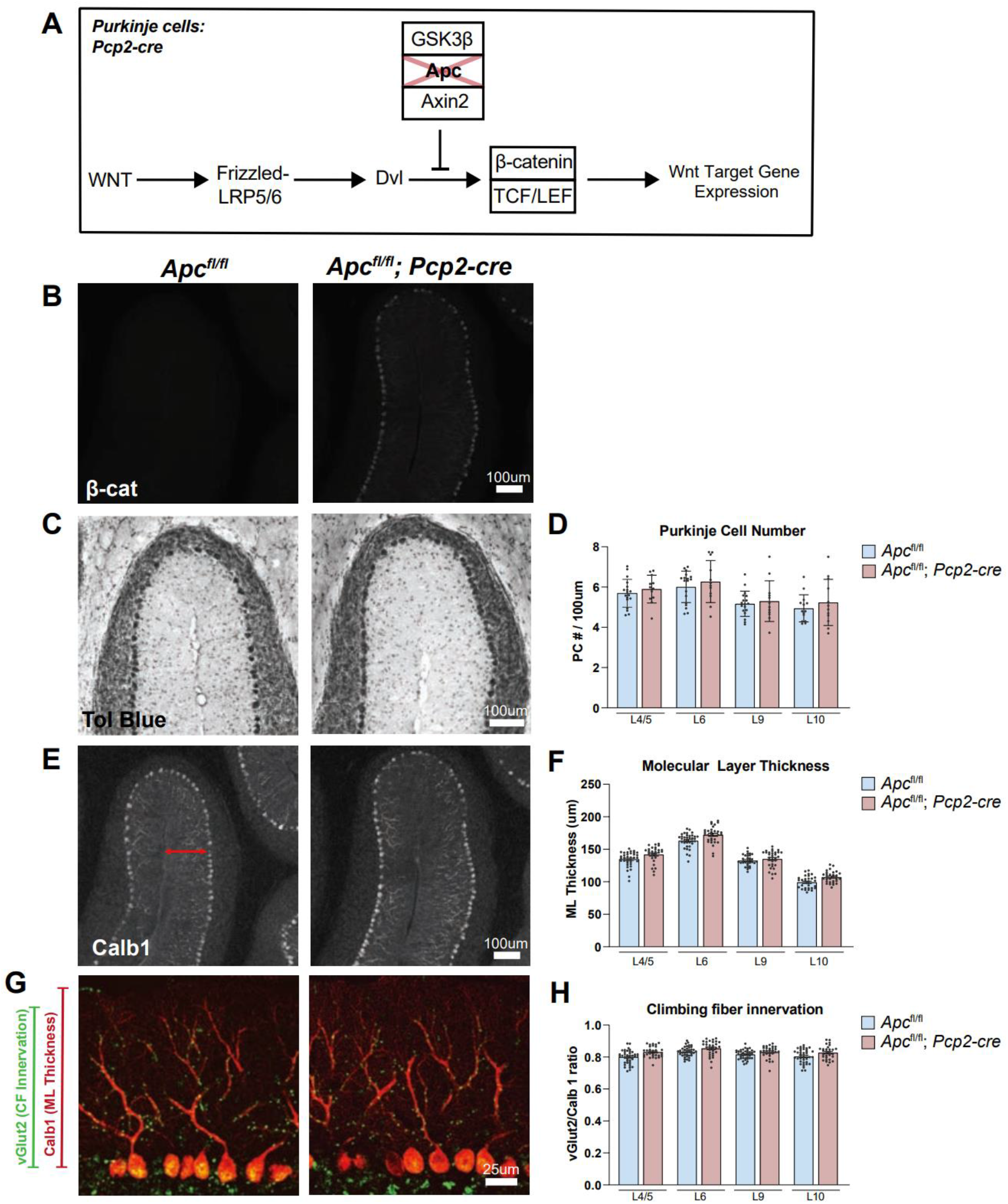
Ectopic activation of Wnt-β-catenin signaling in PCs does not induce SCA1-like phenotypes. **A,** Schematic of Wnt-β-catenin signaling activation in PCs by conditional *Apc* deletion. **B,** Representative images of β-catenin staining in cerebellar lobule 5, showing upregulation of β-catenin intensity in *Apc* PC cKO (*Apc*^fl/fl^; *Pcp2*-cre) mice compared to controls (*Apc*^fl/fl^). **C-H,** Representative images of toluidine blue **(C)**, Calb1 **(E)**, and Calb1 and vGlut2 **(G)** staining of cerebellar lobule 5 in 1 year-old control and *Apc* PC cKO mice, quantified in **(D,F,H)**. Scale bars 100μm **(C,E)**, and 25μm **(G)**. **D,F,H,** Quantifications of PC number per 100μm **(D),** molecular layer thickness in μm **(F)**, and climbing fiber innervation, as ratio of vGlut2 / molecular layer thickness **(H)** in lobules 4/5, 6, 9 and 10 of 1 year-old *Apc* PC cKO and control mice, (*Apc* PC cKO n=5, control n=4). Points in bar plots indicate measurements per image.

We next sought to determine whether activation of Wnt-β-catenin signaling in PCs of WT mice alone was sufficient to induce SCA1-like phenotypes. To activate Wnt-β-catenin signaling in PCs, we generated *Apc* PC cKO (*Apc*^fl/fl^; *Pcp2-cre*) mice in which the *Apc* gene, encoding a key component of the inhibitory destruction complex, was specifically deleted in PCs (Figure 5A). We confirmed elevated β-catenin levels in the cerebellum of *Apc* PC cKO mice by immunostaining (Figure 5B). We observed no significant changes in PC number (Figure 5C,D), molecular layer thickness (Figure 5E,F), or climbing fiber innervation (Figure 5G,H) in 1-year-old *Apc* PC cKO mice compared to controls, demonstrating that activating Wnt-β-catenin signaling specifically in PCs alone is not sufficient to induce SCA1-like pathology. Together, these data suggest that elevated Wnt-β-catenin signaling in PCs observed in SCA1 does not have a profound impact on cerebellar health or disease progression.

### Activation of Wnt-β-catenin signaling in BG induces BG mislocalization and gliosis

As described earlier, we observed enhanced Wnt-β-catenin signaling not only in PCs, but in multiple cerebellar cell types in SCA1 KI and SCA1 Tg [63Q] mice (Figures 1D-H, 3E-H). Because enhanced Wnt-β-catenin signaling in PCs alone was not sufficient to drive SCA1-like phenotypes (Figure 5), and genetically reducing Wnt signaling in PCs was not able to rescue SCA1 phenotypes (Figures 4, S2), we next investigated the impact of elevated Wnt-β-catenin signaling in other cell types of the cerebellum. We were particularly interested in understanding the impact of Wnt-β-catenin signaling in BG, a specialized unipolar astrocyte population exclusive to the cerebellum, for several reasons. First, BG closely associate with PCs and aid in PC synapse function and maintenance in adulthood through glutamate recycling and synaptic finetuning^34, 35^. Second, BG-specific expression of polyQ-expanded ataxin-7, the disease-causing protein for SCA7, impairs glutamate transport via the reduction of GLAST, which is associated with non-cell autonomous PC loss^36^. Interestingly, BG in a SCA1 mouse model similarly display a reduction of GLAST^37^, suggesting dysfunction of BG may contribute to eventual PC loss seen at late stages in SCA1. Finally, previous studies have shown that Wnt-β-catenin signaling activation in BG through *Apc* deletion is sufficient to cause neuronal loss in adult mice and cerebellar degeneration in a non-cell autonomous manner^22^.

First, to confirm that polyQ-expanded mutant ataxin-1 expression in PCs can lead to non-cell autonomous Wnt-β-catenin signaling activation in other cell types, we analyzed Wnt reporter activity in BG of SCA1 Tg [63Q] reporter animals at 12 weeks of age (Figure 6A-D). As was the case with PCs, Wnt reporter activity was significantly increased in Sox9-positive BG in the SCA1 Tg [63Q] mouse cerebellum (Figure 6A-D). This confirmed that, in addition to cell autonomous effects, expression of polyQ-expanded ataxin-1 in PCs leads to the enhanced activation of Wnt-β-catenin signaling in local cell types of the SCA1 cerebellum through indirect mechanisms.

**Figure 6.**
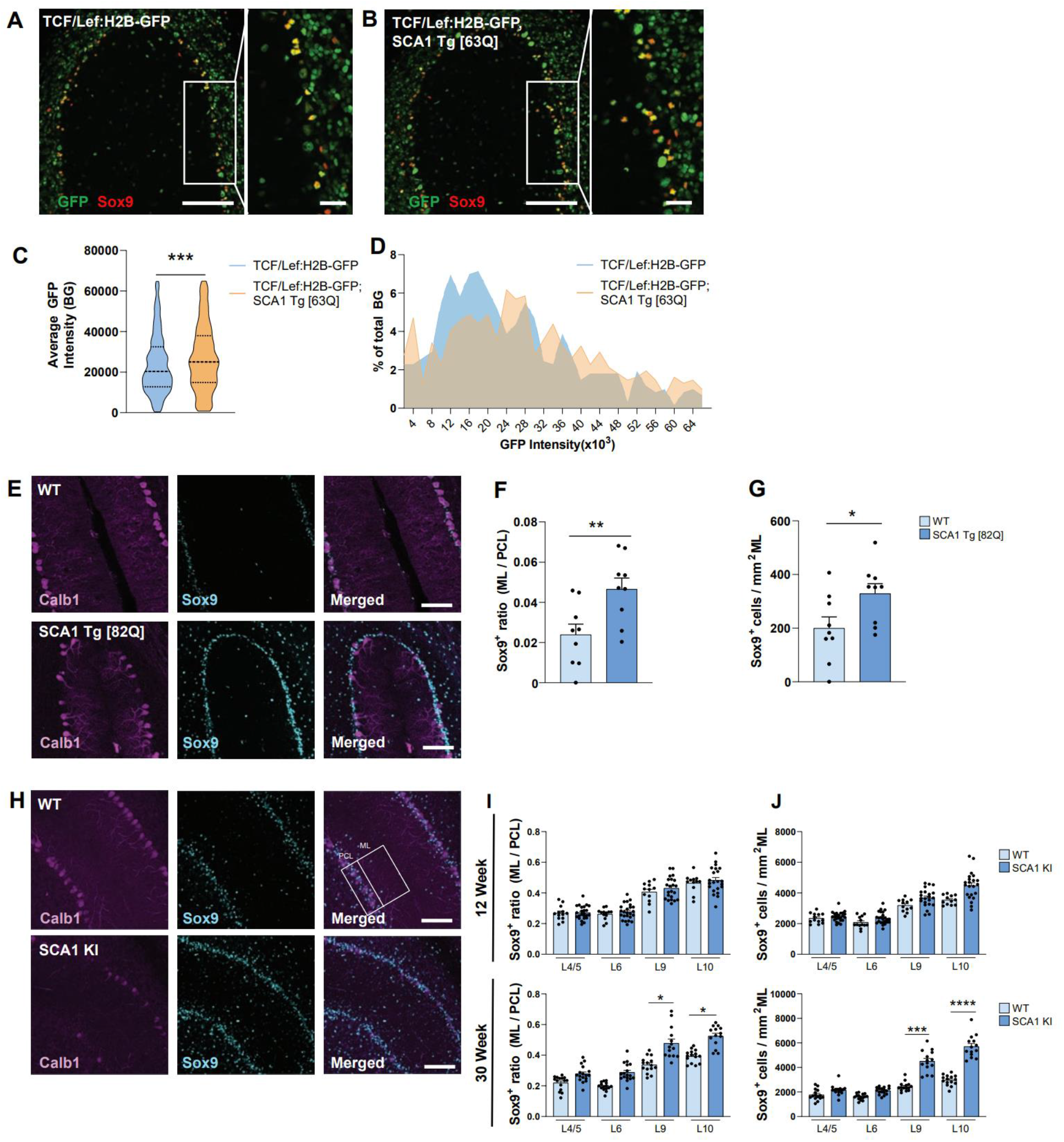
Non-cell autonomous Wnt-β-catenin signaling activation in BG in SCA1 mice and Sox9 mislocalization phenotypes in SCA1. **A-B,** Representative images of 12-week TCF/Lef:H2B-GFP **(A)** and TCF/Lef:H2B-GFP; SCA1 Tg [63Q] **(B)** mouse cerebellar lobule 5, stained with GFP and Sox9. Scale bar 100μm, inset 25μm. **C,** Quantification of intensity of Wnt-β-catenin signaling activity in BG, as average GFP intensity in all Sox9^+^ cells **(C),** and as percentage of total BG counted binned by GFP intensity **(D)**. **E,** Representative images of Calb1^+^ PCs and Sox9^+^ BG in 20-week SCA1 Tg [82Q] and WT cerebellar lobule 4/5. Scale bar 100μm. **F,G,** Quantification of ratio of Sox9^+^ BG in ML/PCL **(F)**, and number of Sox9^+^ BG in ML **(G)** in 20-week cerebellar lobules L4/5 SCA1 Tg [82Q] and WT controls (n=3 per genotype). **H,** Representative images of immunostaining for Calb1^+^ PCs, and Sox9^+^ BG in 30-week WT and SCA1 KI cerebellar lobule 9. Scale bar 100μm. **I,J,** Quantification of ratio of Sox9^+^ BG in ML/PCL **(I)**, and number of Sox9^+^ BG in ML **(J)** in 12 week (top row) and 30 week (bottom row) cerebellar lobules L4/5, L6, L9, and L10 for SCA1 KI and WT controls, WT 12-week n=3, WT 30-week n=3, SCA1 KI 12-week n=4, SCA1 KI 30-week n=5. *P<0.05, **P<0.01, ***P<0.001, ****P<0.0001, by student’s *t*-test (**C**,**F**,**G**), and by one-way ANOVA with Tukey’s post-hoc analysis (**I**,**J**).

We next examined the impact of enhanced Wnt-β-catenin signaling activation in BG. Activation of Wnt-β-catenin signaling in astrocytes (AS) and BG populations through conditional deletion of *Apc* in Gfap-positive cells (*Apc* AS cKO; *Apc*^fl/fl^; *mGFAP*-cre, Figure S3A) was sufficient to increase β-catenin levels and induce severe astrogliosis (Figure S3B,C) and a reduction in body weight in *Apc* AS cKO mice compared to controls (Figure S3D), similar to what is observed in SCA1 mice^38^. Furthermore, BG cell bodies were aberrantly mislocalized from the PC layer (PCL) to the molecular layer (ML) in 4-week-old *Apc* AS cKO mice (Figure S3E-I), consistent with previous studies in which Wnt-β-catenin signaling was activated in BG^22^.

Because Wnt-β-catenin signaling is activated in BG of SCA1 mice (Figure 6A-D) and genetic activation of Wnt-β-catenin signaling in BG is sufficient to induce BG mislocalization during development (Figure S3)^22^, we next wanted to determine whether similar cellular phenotypes occurred in SCA1 mice, in which Wnt-β-catenin activation occurs in BG weeks after the cerebellar cytoarchitecture is established. To this end, we performed immunohistochemical staining for PCs (Calb1^+^ cells) and BG (Sox9^+^ cells) and quantified the number of BG in the PCL and ML (Figures 6E-J, S4). Under physiological conditions, the majority of BG are expected to closely localize to the cell bodies of PCs in the PCL as a monolayer, which was observed in WT animals (Figure 6E-J). Similar to the *Apc* AS cKO mice (Figure S3E-I), we observed a significant increase in the number of heterotopic BG in the ML, as well as significant increases in the ratio of BG cells in the ML/PCL, in 20-week-old SCA1 Tg [82Q] mice (Figure 6E-G) and 30-week-old SCA1 KI mice (Figure 6H-J). Interestingly, no significant differences were detected at 12 weeks in SCA1 KI mice (Figure 6I-J, top row), suggesting BG mislocalization occurs in an age-dependent manner. Considering the important roles of BG in PC maintenance and synapse function^39–43^ and that *Apc* deletion in BG alone has been shown to be sufficient to induce cerebellar degeneration and PC loss^22^, these data suggest that non-cell autonomous activation of Wnt-β-catenin signaling in BG in SCA1 results in disrupted BG to PC interactions that may contribute to cerebellar circuit dysfunction and disease pathogenesis.

## Discussion

The cerebellar cortex circuit is comprised of multiple cell types that intricately interact to produce a coordinated output to the deep cerebellar nuclei via PCs, a population of neurons unique to the cerebellum that undergoes degeneration in many SCAs^2, 44^. How individual cell types contribute to cerebellar dysfunction and PC degeneration in SCAs remains an outstanding question. In this study, we found that ataxin-1, the protein mutated in SCA1, positively regulates Wnt-β-catenin signaling in a polyQ-dependent manner and that canonical Wnt-β-catenin signaling is activated in several SCA1 mouse models in an age-dependent manner in multiple cerebellar cell types, including PCs and BG. To determine the impact of this increased signaling activity in different cell types, we used cell-type specific mouse models to activate or inhibit Wnt-β-catenin signaling in PCs and BG and found that the impact of Wnt-β-catenin signaling in the adult cerebellum was greater in BG rather than PCs, highlighting the importance of glia in SCA1 pathology and disease progression.

We also provide a novel mechanism through which ataxin-1 positively regulates Wnt-β-catenin signaling activity. We found that ataxin-1 enhances Wnt-β-catenin signaling activity at multiple levels of the pathway, and that this is dependent on the presence of pathogenic features of ataxin-1, including polyQ expansion, serine 776-phosphorylation, and nuclear localization. This provides a cell autonomous mechanism for the enhanced activation of Wnt-β-catenin signaling in SCA1 PCs and presumably in other cell types expressing ataxin-1, potentially through direct interaction with TCF/LEF family members. Our studies also provide a potential non-cell autonomous mechanism in which elevated expression of Wnt ligands in any given cell type could activate Wnt-β-catenin signaling in surrounding cell types. Enhanced Wnt ligand secretion may also explain the dysregulation of genes associated with planar cell polarity (PCP) signaling, a noncanonical Wnt signaling pathway, observed in SCA1 in previous studies^8^.Although expression of mutant ataxin-1 in the cerebellum is primarily limited to PCs in the transgenic animal models used here, the cellular source of enhanced Wnt ligand production is unclear. While it is possible that PCs expressing polyQ-expanded ataxin-1 may cell autonomously increase Wnt ligand production, it is also possible that PC dysfunction may non-cell autonomously stimulate other local cell types of the cerebellar cortex to increase production of Wnt ligands. Furthermore, although increased secretion of Wnt ligands can contribute to local Wnt activation in adjacent cell types, it is possible that non-cell autonomous Wnt activity can also be influenced by altered inflammatory states of the various glial populations of the cerebellum, in which changes in intracellular signaling pathways that reciprocally regulate Wnt-β-catenin signaling, such as NF-κB, have been reported^38, 45, 46^. However, the degree to which activation of BG contributes to enhanced Wnt activity in SCA1 independent of extracellular Wnt ligands, remains to be determined.

We observed enhanced Wnt-β-catenin signaling in multiple cell types, besides PCs and BG, in our SCA1 mouse models in an age-dependent manner compared to controls. Future studies to investigate which other cell types exhibit enhanced Wnt-β-catenin signaling in SCA1 mice are needed. Furthermore, whether this activation of Wnt-β-catenin signaling in those cell types is protective, toxic, compensatory, or secondary to SCA1 disease progression requires further investigation. Interestingly, Wnt signaling has been implicated in other neurodegenerative diseases, including Alzheimer’s disease (AD)^47–51^, amyotrophic lateral sclerosis (ALS)^52, 53^, and Huntington’s Disease (HD)^54^. Previous studies of the impact of Wnt-β-catenin signaling in AD suggest enhanced signaling may be protective against the toxicity of Aβ peptides, including synapse loss^55^. In contrast, enhanced β-catenin levels are thought to be toxic in HD^54^. Our studies here in SCA1 suggest that activation of Wnt-β-catenin signaling impacts unique cell populations in the cerebellum differently, and highlight the importance of examining signaling pathways in the specific cellular contexts in which they may be affected. Our data indicate that while activation of Wnt-β-catenin in PCs does not significantly contribute to SCA1 phenotypes, activation of Wnt-β-catenin signaling in BG is detrimental and sufficient to cause gliosis and BG mislocalization in *Apc* AS cKO mice. The exact cellular mechanisms through which BG impact PC health and SCA1 pathology are still unclear; however, these findings underscore the complex cell-cell interactions underlying SCA1 pathogenesis and bring attention to the role of BG in PC survival and in SCA1 disease progression.

## Materials and Methods

### Animal husbandry

All animal care procedures were approved by the Yale University Institutional Animal Care and Use Committee. Mice were kept in a 12 hour light/dark cycle with standard chow and *ad libitum* access to water. A combination of males and females were used for all experiments. Three mouse models of SCA1 were utilized; a SCA1 knock-in (SCA1 KI; *Atxn1*^154Q/+^)^26^ strain which expresses mutant ataxin-1 with 154 glutamine repeats under its endogenous promoter, a SCA1 transgenic (SCA1 Tg [82Q]; *Pcp2-ATXN1*^82Q/+^)^33^ line in which mutant ataxin-1 with 82 glutamine repeats is overexpressed under the PC-specific *Pcp2* promoter, and a SCA1 transgenic (SCA1 Tg [63Q]; *Pcp2-ATXN1*^63Q/+^) line, in which the polyQ tract length was reduced to 63 glutamine repeats due to a germline contraction event. These lines were crossed with a Wnt-β-catenin reporter line (TCF/Lef:H2B-GFP)^27^, in which Wnt-β-catenin activity drives GFP expression in cells. Additionally, the SCA1 Tg [63Q] and SCA1 Tg [82Q] lines were crossed with *Ctnnb1* PC cKO (*Ctnnb1*^fl/fl^; *Pcp2*-cre), *Apc* PC cKO (*Apc*^fl/fl^; *Pcp2*-cre), and *Apc* AS cKO (*Apc*^fl/fl^; *mGFAP*-cre) lines, in which Wnt-β-catenin signaling is activated or inhibited in PC (under the PC-specific *Pcp2* promoter) or astrocyte (under the astrocyte-specific *mGFAP* promoter) populations specifically. *Ctnnb1*^fl/fl^ (Jackson Laboratory Stock No. 004152), TCF/Lef:H2B-GFP (Jackson Laboratory Stock No. 013752), *mGFAP*-cre (Jackson Laboratory Stock No. 012886), and *Apc*^fl/fl^ (EMMA mouse repository, Stock No. EM:05566) strains were purchased commercially. SCA1 Tg [63Q] mice were originally maintained on a FVB background and backcrossed onto C57BL/6J for three generations before analyses. All other mice and littermates were maintained on a C57BL/6J background.

### Protein extraction and western blot analysis

Mouse cerebellar tissue was homogenized in Triple lysis buffer (0.5% NP-40, 0.5% Triton X-100, 0.1% SDS, 20 mM Tris-HCl (pH 8.0), 180 mM NaCl, 1 mM EDTA and Roche complete protease inhibitor cocktail and PhosStop protease inhibitors) by dounce homogenization on ice. Samples were then sonicated to ensure breakdown of protein aggregates before rotation at 4°C for 10 minutes and centrifugation for 10 minutes at 13,000 rpm at 4°C. Total protein concentration of the supernatant was quantified using a BCA assay (ThermoFisher 23225) and equal protein amounts were boiled at 95°C for 10 minutes prior to being run on a gel (BioRad) at 120V. Proteins from gels were transferred for one hour at 100V at 4°C onto 0.45μm nitrocellulose membranes. Membranes were washed with TBST (Tris-buffered saline, 0.1% Tween-20) three times for 10 minutes each, followed by one hour of blocking in 5% non-fat dry milk in TBST at room temperature. Membranes were then incubated with primary antibody in 5% non-fat dry milk in TBST at 4°C overnight. The following day, membranes were washed with TBST three times for 10 minutes each, followed by incubation in either sheep anti-mouse or donkey anti-rabbit IgG-conjugated with horseradish peroxidase (HRP) (Millipore Sigma, GENA931, GENA934, 1:4,000) in TBST at room temperature for two hours. Membranes were then washed with TBST three times for 10 minutes and developed using SuperSignal West Pico Plus Chemiluminescent substrate (Pierce, Cat. 34580) and visualized using a KwikQuant Imager (Kindle Biosciences). Images were quantified using ImageJ. The following primary antibodies were used: Mouse anti-Gapdh (Sigma G8795, 1:10,000), mouse anti-β-catenin (BD 610154, 1:20,000), mouse anti-active β-catenin (Millipore 06-665 1:500), rabbit anti-ataxin-1 (11750, 1:1,000), mouse anti-T7 (Novagen 69522, 1:10,000), rabbit anti-HA (Abcam ab9110, 1:5,000), mouse anti-c-Myc (Sigma M5546 clone 9e10, 1:1,000), and rabbit anti-GST (Sigma G7781, 1:3,000).

### RNA extraction and RT-qPCR

RNA was extracted from frozen mouse cerebellar tissue using the Qiagen RNeasy Mini Kit, following the manufacturer’s instructions. cDNA was synthesized using oligo-dT primers and iScript cDNA synthesis kit (BioRad, 1708891). Reverse transcription-quantitative Polymerase Chain Reaction (RT-qPCR) was performed using TaqMan probes with iTaq Universal Probe Supermix on a C1000 Thermal Cycler (BioRad). The following TaqMan (Applied Biosystems) probes were used: *Gapdh* (4352661, Mm99999915_g1), *Actb* (4352933E, Mm00607939_s1), *Hprt* (4331182, Mm03024075_m1), *Ccnd1* (4331182, Mm00432359_m1), *Myc* (4331182, Mm00487804_m1), *Axin2* (4331182, Mm00443610_m1). Custom TaqMan array plates (Thermofisher, 4413261) were used for the Wnt ligand screen, with the following TaqMan probes: *Actb* (Mm00607939_s1), *Gapdh* (Mm99999915_g1), *Hprt* (Mm00446968_m1), *Wnt1* (Mm01300555_g1), *Wnt2* (Mm00470018_m1), *Wnt2b* (Mm00437330_m1), *Wnt3* (Mm00437336_m1), *Wnt4* (Mm01194003_m1), *Wnt5a* (Mm00437347_m1), *Wnt6* (Mm00437353_m1), *Wnt7a* (Mm00437356_m1), *Wnt8b* (Mm00442108_g1), *Wnt10a* (Mm00437325_m1), *Wnt10b* (Mm00442104_m1), *Wnt11* (Mm00437327_g1). All samples were loaded in triplicate. Target gene expression was normalized to housekeeping genes (*Actb*, *Gapdh*, and *Hprt*) using BioRad CFX manager software and plotted relative to mean expression of WT littermate controls.

### Immunofluorescence staining

Mice were anesthetized prior to intracardial perfusion with phosphate buffered saline (PBS) and 4% paraformaldehyde (PFA). Brains were post-fixed overnight in 4% PFA before incubation in 20% and 30% sucrose in PBS. Samples were frozen in OCT compound (VWR, 4538) and sliced into 30μm sections on a cryostat (Leica). Free-floating sections were washed in PBS and PBS with 0.1% Triton-X before incubation in 5% normal goat serum (Jackson Labs, 005-000-121) at room temperature. Upon usage, antigen retrieval with 10mM citric acid for 30 minutes was used before wash and incubation in primary antibody. Primary antibody incubation was carried out at 4°C overnight with the following antibodies: mouse anti-Calbindin-D-28K (Sigma, C9848, 1:1000), rabbit anti-vGlut2 (Synaptic Systems, 135402, 1:500), rabbit anti-β-Catenin (Sigma, AV14001,1:1000), mouse anti-β-catenin (BD Transduction Laboratories, 610154, 1:1000), chicken anti-Gfap (Abcam, ab4674, 1:1000), rabbit anti-Iba1 (Wako, 019-19741, 1:500), rabbit anti-Sox9 (1:500), and rabbit anti-GFP (Abcam, ab290, 1:4000). Sections were washed in PBS with 0.1% Triton-X before incubation in secondary antibody (Invitrogen AlexaFluor, 1:500) then washed and mounted onto slides and coverslipped with Vectashield mounting media and DAPI (Vector Laboratories, H-1500). Fluorescent images were acquired on a Zeiss LSM800 or LSM880 confocal microscope, using the same microscope and settings across similar experiments. Between 3-6 brain sections were imaged and quantified for each mouse.

### Fluorescent image quantification

ImageJ (National Institutes of Health) was used for all image processing and quantification. For fluorescent intensity quantification, image z-stacks were flattened to maximum intensity z-projection, converted to 8-bit images, and thresholded using identical parameters across all images. ROI manager was used to measure equal areas across images. For GFP fluorescent intensity quantification in PCs or BG, calbindin (for PCs) or Sox9 (for BG) images were thresholded, and overlaid onto the GFP images. The mean gray intensity value was recorded. To measure the molecular layer thickness, the lengths of the PC dendrites were measured at six locations in the image, from the tip of the PC soma until the end of the molecular layer approximately 300μm from the tip of the specified cerebellar lobule. For calbindin intensity across the molecular layer, a similar methodology was employed as previously described^26^. Briefly, the z-stack was flattened in ImageJ using the average intensity z-projection, and the intensity per pixel plotted using plot profile. Three PCs approximately 300μm from the tip of the lobule per section were used for quantification, and the average intensity value for 1μm increments plotted for each animal. For assessing the innervation of inferior olive climbing fiber vGlut2 puncta along PC dendrites, a z-stack corresponding to a 5μm depth was flattened in ImageJ using the maximum intensity z-projection. The length of vGlut2 innervation relative to the length of the molecular layer was measured at three separate locations along the lobule, similar to the methodology for assessing calbindin intensity. For assessing Sox9-positive BG distribution, image z-stacks were flattened to maximum intensity z-projection, converted to 8-bit images, and thresholded using equal parameters across all images. ROI manager was used to measure equal areas across images for the PC layer and molecular layer. The watershed function was utilized to split clumped nuclei, and only BG with nuclei greater than 3μm^2^ were analyzed. For all quantifications, 3-6 images were quantified per mouse, and data was plotted using GraphPad Prism.

### Toluidine blue staining and quantification

Perfused brain sections 30μm thick were mounted onto slides, outlined with a Pap pen, and washed with PBS before incubating in 2% toluidine blue in PBS for approximately 5 minutes. Sections were monitored under a dissection microscope until PCs were adequately labeled. Slides were then washed in PBS until residual stain was removed, then allowed to dry before coverslipping with Permount. Sections were imaged on an Olympus microscope and PC counts were quantified by a blinded observer in ImageJ. The total number of PCs across a 200μm distance was quantified.

### Luciferase reporter assay

HeLa cells were plated at 7.5×10^4^ cells per well 24 hours prior to transfection. Cells were transfected with 10ng TOP or FOP flash, 1ng Renilla, and 100ng of gene of interest. For Wnt3a conditioned media, 24 hours later media was changed and incubated overnight. Luciferase activity was measured 48 hours post-transfection (Dual luciferase, Promega).

### Co-affinity purification

HeLa cells were plated at 0.5-1×10^6^ cells per well 24 hours prior to transfection, maintained in a 37°C, 5.5% CO_2_ incubator. When 70-80% confluent, cells were transfected with 0.5μg of gene of interest, 1μg of GST-ATXN1 plasmid, 4.5μg polyethylenimine (PEI), and Opti-MEM. 48 hours post-transfection, cells were transferred into 1 mL PBS and centrifuged for one minute at 13,000rpm. The cells were resuspended in Triple lysis buffer, and rotated at 4°C for 30min to lyse cells. Cells were centrifuged for 15 minutes at 13,000rpm at 4°C. Supernatant was either kept for crude extract, or transferred to 20μl of washed glutathione-sepharose 4B beads (Millipore Sigma, GE17-0756-01) for affinity purification. Affinity purification samples were rotated overnight at 4°C and washed 5 times in lysis buffer. All samples were diluted in 4x buffer with BME, and boiled for 10 minutes at 95°C prior to loading gel. Western blots were run as described above. The following primary antibodies were used: T7 (Novagen 69522, 1:10,000), HA (Abcam ab9110, 1:5,000), c-Myc (Sigma M5546 clone 9e10, 1:1,000), β-catenin (BD 610154, 1:20,000), and GST (Sigma G7781, 1:3,000). To visualize GST signal, membranes were stripped with 10mM sodium azide in TBST for one hour, washed three times with TBST for 10 minutes each, and blocked with 5% non-fat milk in TBST for 1 hour, prior to incubation with GST primary antibody overnight at 4°C. Subsequent steps were performed as described above.

### Statistical analysis

All statistical analyses were performed in GraphPad Prism. Data are shown as mean ± SEM. To determine statistical significance, either two-tailed unpaired Student’s *t* tests (when comparing two experimental groups) or one-way analysis of variance (ANOVA) with Tukey’s post-hoc analysis (when comparing more than two experimental groups) were performed, with a significance cutoff of *P* < 0.05.

## Supporting information

Luttik et al. - Supplementary Information

## Acknowledgements

We thank all members from the Lim laboratory for technical support, thoughtful discussion, comments and critiques for this project. This work was supported by National Institute of Health grants NS083706 (J.L.), NS088321 (J.L.), MH119803 (J.L.), AG066447 (J.L.), and T32 NS041228 (K.L., L.T.).

## References

1. Orr, H.T., et al. Expansion of an unstable trinucleotide CAG repeat in spinocerebellar ataxia type 1. Nat Genet 4, 221–226 (1993).

2. Orr, H.T. & Zoghbi, H.Y. SCA1 molecular genetics: a history of a 13 year collaboration against glutamines. Hum Mol Genet 10, 2307–2311 (2001).

3. Driessen, T.M., Lee, P.J. & Lim, J. Molecular pathway analysis towards understanding tissue vulnerability in spinocerebellar ataxia type 1. Elife 7 (2018).

4. Serra, H.G., et al. RORalpha-mediated Purkinje cell development determines disease severity in adult SCA1 mice. Cell 127, 697–708 (2006).

5. Edamakanti, C.R., Do, J., Didonna, A., Martina, M. & Opal, P. Mutant ataxin1 disrupts cerebellar development in spinocerebellar ataxia type 1. J Clin Invest 128, 2252–2265 (2018).

6. Lin, X., Antalffy, B., Kang, D., Orr, H.T. & Zoghbi, H.Y. Polyglutamine expansion down-regulates specific neuronal genes before pathologic changes in SCA1. Nat Neurosci 3, 157–163 (2000).

7. Ruegsegger, C., et al. Impaired mTORC1-Dependent Expression of Homer-3 Influences SCA1 Pathophysiology. Neuron 89, 129–146 (2016).

8. Ingram, M., et al. Cerebellar Transcriptome Profiles of ATXN1 Transgenic Mice Reveal SCA1 Disease Progression and Protection Pathways. Neuron 89, 1194–1207 (2016).

9. Lein E.S, et al. Genome-wide atlas of gene expression in the adult mouse brain. Nature 445, 168–176 (2007).

10. Servadio, A., et al. Expression analysis of the ataxin-1 protein in tissues from normal and spinocerebellar ataxia type 1 individuals. Nat Genet 10, 94–98 (1995).

11. Zhang, Y., et al. An RNA-sequencing transcriptome and splicing database of glia, neurons, and vascular cells of the cerebral cortex. J Neurosci 34, 11929–11947 (2014).

12. Thomas, K.R., Musci, T.S., Neumann, P.E. & Capecchi, M.R. Swaying is a mutant allele of the proto-oncogene Wnt-1. Cell 67, 969–976 (1991).

13. Goold, R., et al. Down-regulation of the dopamine receptor D2 in mice lacking ataxin 1. Hum Mol Genet 16, 2122–2134 (2007).

14. Zeng, L., et al. Loss of the Spinocerebellar Ataxia type 3 disease protein ATXN3 alters transcription of multiple signal transduction pathways. PLoS ONE 13 (2018).

15. Komiya, Y. & Habas, R. Wnt signal transduction pathways. Organogenesis 4, 68–75 (2008).

16. Patapoutian, A. & Reichardt, L.F. Roles of Wnt proteins in neural development and maintenance. Curr Opin Neurobiol 10, 392–399 (2000).

17. Brault, V., et al. Inactivation of the beta-catenin gene by Wnt1-Cre-mediated deletion results in dramatic brain malformation and failure of craniofacial development. Development 128, 1253–1264 (2001).

18. Selvadurai, H.J. & Mason, J.O. Activation of Wnt/beta-catenin signalling affects differentiation of cells arising from the cerebellar ventricular zone. PLoS One 7, e42572 (2012).

19. Hall, A.C., Lucas, F.R. & Salinas, P.C. Axonal remodeling and synaptic differentiation in the cerebellum is regulated by WNT-7a signaling. Cell 100, 525–535 (2000).

20. Pei, Y., et al. WNT signaling increases proliferation and impairs differentiation of stem cells in the developing cerebellum. Development 139, 1724–1733 (2012).

21. Lorenz, A., et al. Severe alterations of cerebellar cortical development after constitutive activation of Wnt signaling in granule neuron precursors. Mol Cell Biol 31, 3326–3338 (2011).

22. Wang, X., Imura, T., Sofroniew, M.V. & Fushiki, S. Loss of adenomatous polyposis coli in Bergmann glia disrupts their unique architecture and leads to cell nonautonomous neurodegeneration of cerebellar Purkinje neurons. Glia 59, 857–868 (2011).

23. Schuller, U. & Rowitch, D.H. Beta-catenin function is required for cerebellar morphogenesis. Brain Res 1140, 161–169 (2007).

24. Brakeman, J.S., Gu, S.H., Wang, X.B., Dolin, G. & Baraban, J.M. Neuronal localization of the Adenomatous polyposis coli tumor suppressor protein. Neuroscience 91, 661–672 (1999).

25. Coyle-Rink, J., Valle, D., L, S. & T, K. Developmental expression of Wnt signaling factors in mouse brain. Cancer Biol Ther 1, 640–645 (2002).

26. Watase, K., et al. A long CAG repeat in the mouse Sca1 locus replicates SCA1 features and reveals the impact of protein solubility on selective neurodegeneration. Neuron 34, 905–919 (2002).

27. Ferrer-Vaquer, A., et al. A sensitive and bright single-cell resolution live imaging reporter of Wnt/B-catenin signaling in the mouse. BMC Dev Biol 10, 121–121 (2010).

28. Friedrich, J., et al. Antisense oligonucleotide-mediated ataxin-1 reduction prolongs survival in SCA1 mice and reveals disease-associated transcriptome profiles. JCI Insight 3 (2018).

29. Veeman, M.T., Slusarski, D.C., Kaykas, A., Louie, S.H. & Moon, R.T. Zebrafish prickle, a modulator of noncanonical Wnt/Fz signaling, regulates gastrulation movements. Curr Biol 13, 680–685 (2003).

30. MacDonald, B.T., Tamai, K. & He, X. Wnt/beta-catenin signaling: components, mechanisms, and diseases. Dev Cell 17, 9–26 (2009).

31. Tejwani, L. & Lim, J. Pathogenic mechanisms underlying spinocerebellar ataxia type 1. Cell Mol Life Sci (2020).

32. Zoghbi, H.Y. & Orr, H.T. Pathogenic mechanisms of a polyglutamine-mediated neurodegenerative disease, spinocerebellar ataxia type 1. J Biol Chem 284, 7425–7429 (2009).

33. Burright, E.N., et al. SCA1 transgenic mice: a model for neurodegeneration caused by an expanded CAG trinucleotide repeat. Cell 82, 937–948 (1995).

34. Grosche, J., et al. Microdomains for neuron-glia interaction: parallel fiber signaling to Bergmann glial cells. Nat Neurosci 2, 139–143 (1999).

35. Yamada, K., et al. Dynamic transformation of Bergmann glial fibers proceeds in correlation with dendritic outgrowth and synapse formation of cerebellar Purkinje cells. J Comp Neurol 418, 106–120 (2000).

36. Custer, S.K., et al. Bergmann glia expression of polyglutamine-expanded ataxin-7 produces neurodegeneration by impairing glutamate transport. Nature Neuroscience 9, 1302–1311 (2006).

37. Cvetanovic, M. Decreased expression of glutamate transporter GLAST in Bergmann glia is associated with the loss of Purkinje neurons in the spinocerebellar ataxia type 1. Cerebellum 14, 8–11 (2015).

38. Cvetanovic, M., Ingram, M., Orr, H. & Opal, P. Early activation of microglia and astrocytes in mouse models of spinocerebellar ataxia type 1. Neuroscience 289, 289–299 (2015).

39. Buffo, A. & Rossi, F. Origin, lineage and function of cerebellar glia. 42–63 (2013).

40. Bellamy, T.C. Interactions between Purkinje neurones and Bergmann glia. Cerebellum 5, 116–126 (2006).

41. De Zeeuw, C.I. & Hoogland, T.M. Reappraisal of Bergmann glial cells as modulators of cerebellar circuit function. Frontiers in Cellular Neuroscience 9, 1–8 (2015).

42. Sasaki, T., et al. Application of an optogenetic byway for perturbing neuronal activity via glial photostimulation. Proc Natl Acad Sci U S A 109, 20720–20725 (2012).

43. Yamada, K. & Watanabe, M. Cytodifferentiation of Bergmann glia and its relationship with Purkinje cells. Anat Sci Int 77, 94–108 (2002).

44. Robinson, K.J., Watchon, M. & Laird, A.S. Aberrant Cerebellar Circuitry in the Spinocerebellar Ataxias. Front Neurosci 14, 707 (2020).

45. Kim, J.H., Lukowicz, A., Qu, W., Johnson, A. & Cvetanovic, M. Astroglia contribute to the pathogenesis of spinocerebellar ataxia Type 1 (SCA1) in a biphasic, stage-of-disease specific manner. Glia 66, 1972–1987 (2018).

46. Ma, B. & Hottiger, M.O. Crosstalk between Wnt/beta-Catenin and NF-kappaB Signaling Pathway during Inflammation. Front Immunol 7, 378 (2016).

47. Inestrosa, N.C., Varela-Nallar, L., Grabowski, C.P. & Colombres, M. Synaptotoxicity in Alzheimer’s disease: the Wnt signaling pathway as a molecular target. IUBMB Life 59, 316–321 (2007).

48. Toledo, E.M. & Inestrosa, N.C. Activation of Wnt signaling by lithium and rosiglitazone reduced spatial memory impairment and neurodegeneration in brains of an APPswe/PSEN1DeltaE9 mouse model of Alzheimer’s disease. Mol Psychiatry 15, 272–285 (2010).

49. Alvarez, A.R., et al. Wnt-3a overcomes beta-amyloid toxicity in rat hippocampal neurons. Exp Cell Res 297, 186–196 (2004).

50. da Cruz e Silva, O.A., Henriques, A.G., Domingues, S.C. & da Cruz e Silva, E.F. Wnt signalling is a relevant pathway contributing to amyloid beta-peptide-mediated neuropathology in Alzheimer’s disease. CNS Neurol Disord Drug Targets 9, 720–726 (2010).

51. Li, B., et al. WNT5A signaling contributes to Abeta-induced neuroinflammation and neurotoxicity. PLoS One 6, e22920 (2011).

52. Chen, Y., et al. Wnt signaling pathway is involved in the pathogenesis of amyotrophic lateral sclerosis in adult transgenic mice. Neurol Res 34, 390–399 (2012).

53. Chen Y, et al. Activation of the Wnt/beta-catenin signaling pathway is associated with glial proliferation in the adult spinal cord of ALS transgenic mice. Biochem Biophys Res Commun 420, 397–403 (2012).

54. Godin, J.D., Poizat, G., Hickey, M.A., Maschat, F. & Humbert, S. Mutant huntingtin-impaired degradation of beta-catenin causes neurotoxicity in Huntington’s disease. EMBO J 29, 2433–2445 (2010).

55. Inestrosa, N.C. & Varela-Nallar, L. Wnt signaling in the nervous system and in Alzheimer’s disease. Journal of Molecular Cell Biology 6, 64–74 (2014).

