## Supplementary material for "Differential effects of Wnt-β-catenin signaling in Purkinje cells and Bergmann glia in SCA1": Luttik et al. - Supplementary Information

**Affiliations:**
[1]Interdepartmental Neuroscience Program, Yale School of Medicine, New Haven, CT 06510, USA
[2]Department of Neuroscience, Yale School of Medicine, New Haven, CT 06510, USA
[3]Department of Genetics, Yale School of Medicine, New Haven, CT 06510, USA
[4]Program in Cellular Neuroscience, Neurodegeneration and Repair, Yale School of Medicine, New Haven, CT 06510, USA
[5]Yale Stem Cell Center, Yale School of Medicine, New Haven, CT 06510, USA

**Classification:** Biological Sciences, Neuroscience

**Keywords:** spinocerebellar ataxia type 1, SCA1, neurodegeneration, Wnt signaling, Purkinje cells, Bergmann glia

**This PDF file includes:**

Figures S1 to S4

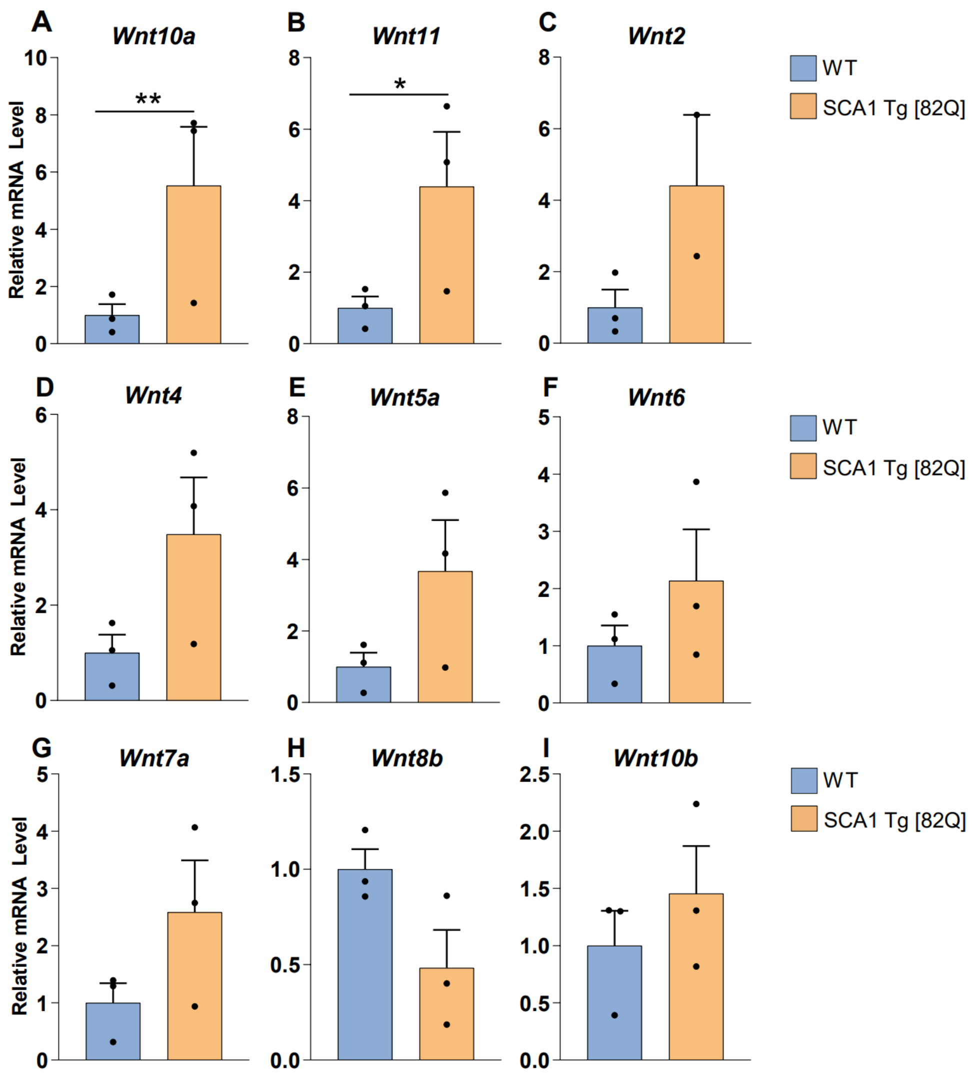


Luttik et al., Figure S1

**Figure S1, *Supplementary to Figure 3;* Expression of secreted Wnt ligands in SCA1 Tg [82Q] mice. A-I,** Relative mRNA expression levels of secreted Wnt ligands *Wnt10a* **(A)**, *Wnt11* **(B)**, *Wnt2* **(C)**, *Wnt4* **(D)**, *Wnt5a* **(E)**, *Wnt6* **(F)**, *Wnt7a* **(G)**, *Wnt8b* **(H)**, and *Wnt10b* **(I)** in 30-week SCA1 Tg [82Q] mouse cerebellum, normalized to WT controls, n=3 animals per genotype. *P<0.05, **P<0.01, by student's *t*-test.

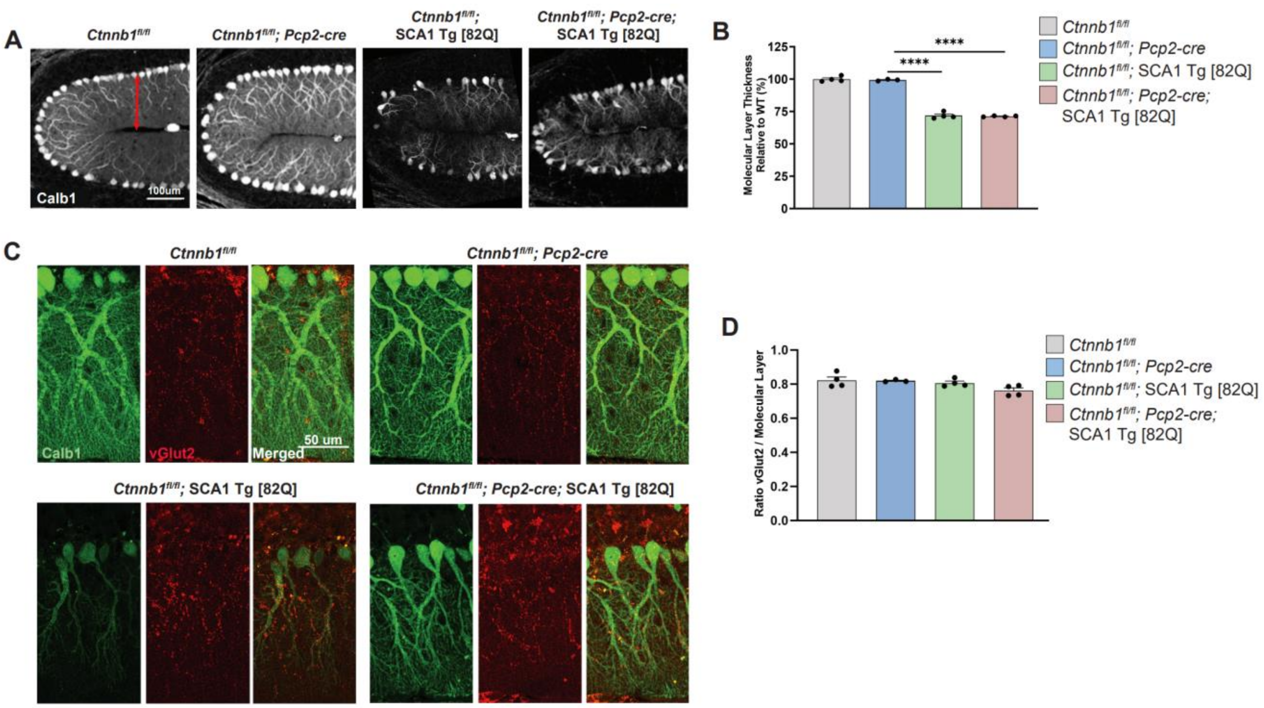


**Luttik et al., Figure S2**

**Figure S2, *Supplementary to Figure 4;* Loss of β-catenin in PCs on SCA1 Tg [82Q] background does not prevent SCA1 Tg [82Q] phenotypes. A,B,** Representative images of Calb1 staining of 21-week cerebellar lobule 4/5 in control and *Ctnnb1* PC cKO (*Ctnnb1*$^{fl/fl}$; *Pcp2-cre*) mice on WT and SCA1 Tg [82Q] backgrounds **(A)**, to measure molecular layer thickness relative to *Ctnnb1*$^{fl/fl}$ controls **(B),** n=4, 3, 4, 4. **C,D,** Representative images of Calb1 and vGlut2 staining of 21-week cerebellar lobule 4/5 in control and *Ctnnb1* PC cKO mice on WT and SCA1 Tg [82Q] backgrounds **(C)**, to measure climbing fiber innervation quantified in **(D),** n=4, 3, 4, 4. ****P<0.0001, by one-way ANOVA with Tukey's post-hoc analysis.

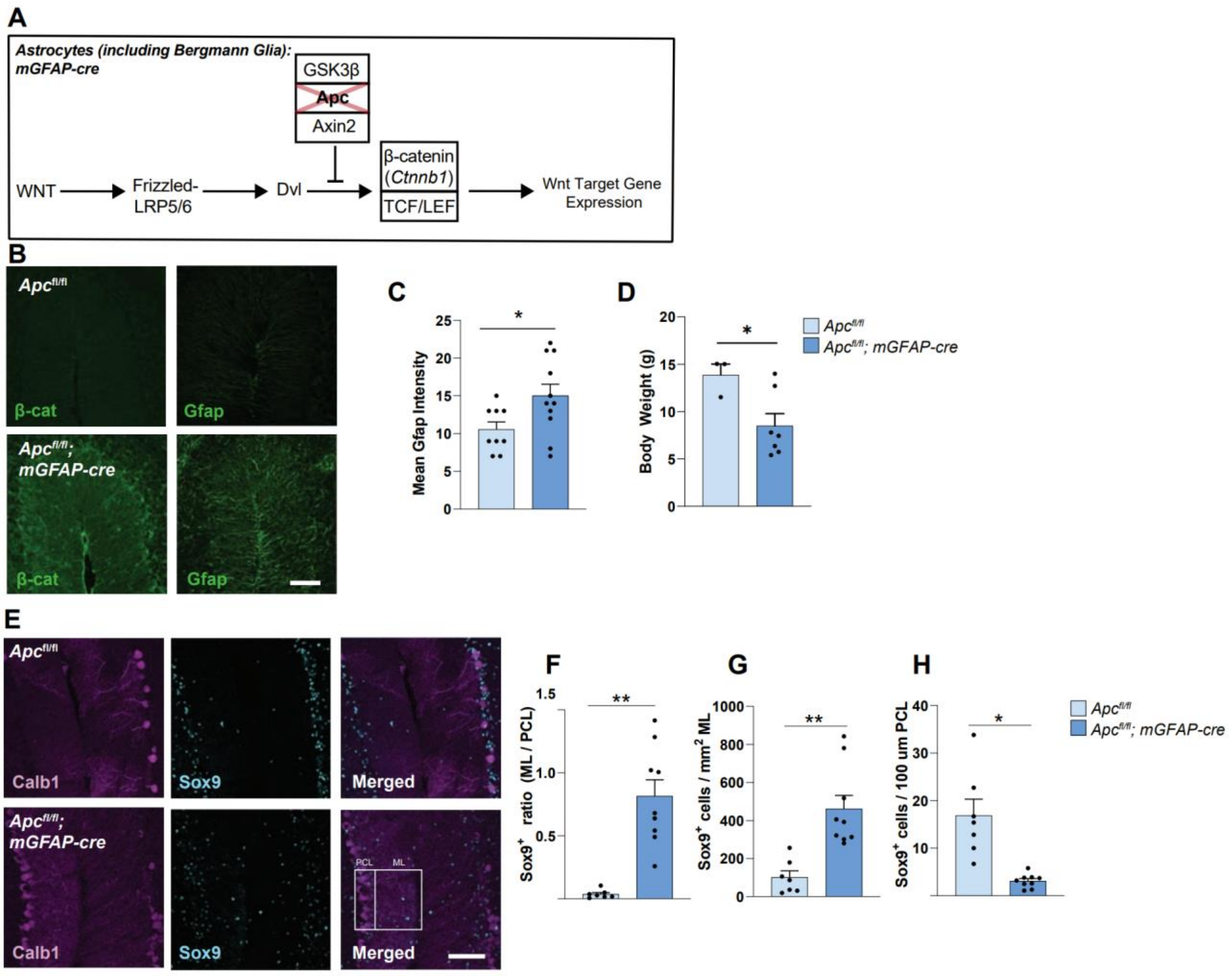


Luttik et al., Figure S3

**Figure S3, *Supplementary to Figure 6;* Ectopic activation of Wnt-β-catenin signaling in BG/astrocyte populations leads to gliosis and BG mislocalization. A,** Schematic of Wnt-β-catenin signaling activation in BG/astrocyte by *Apc* conditional deletion. **B,** Representative images of immunostaining for β-catenin and Gfap in 4-week control (*Apc*$^{fl/fl}$), and *Apc* AS cKO (*Apc*$^{fl/fl}$; *mGFAP*-cre) mice, n=4 per genotype. Scale bar 100μm. **C,** Quantification of mean Gfap intensity in (**B**). **D,** Body weights of control and *Apc* AS cKO mice at 4 weeks. **E,** Representative images of immunostaining for Calb1$^+$ PCs and Sox9$^+$ BG in control and *Apc* AS cKO mice at 4 weeks, n=4 per genotype. Box indicating regions defined as PC layer (PCL) and molecular layer (ML). Scale bar 100μm. **F-I,** Quantification of ratio of Sox9$^+$ BG in ML/PCL **(F)**, and number of Sox9$^+$ BG in ML **(G)**, and PCL **(H)** in 4-week *Apc* AS cKO mice and littermate controls. *P<0.05, **P<0.01, by student's *t*-test.

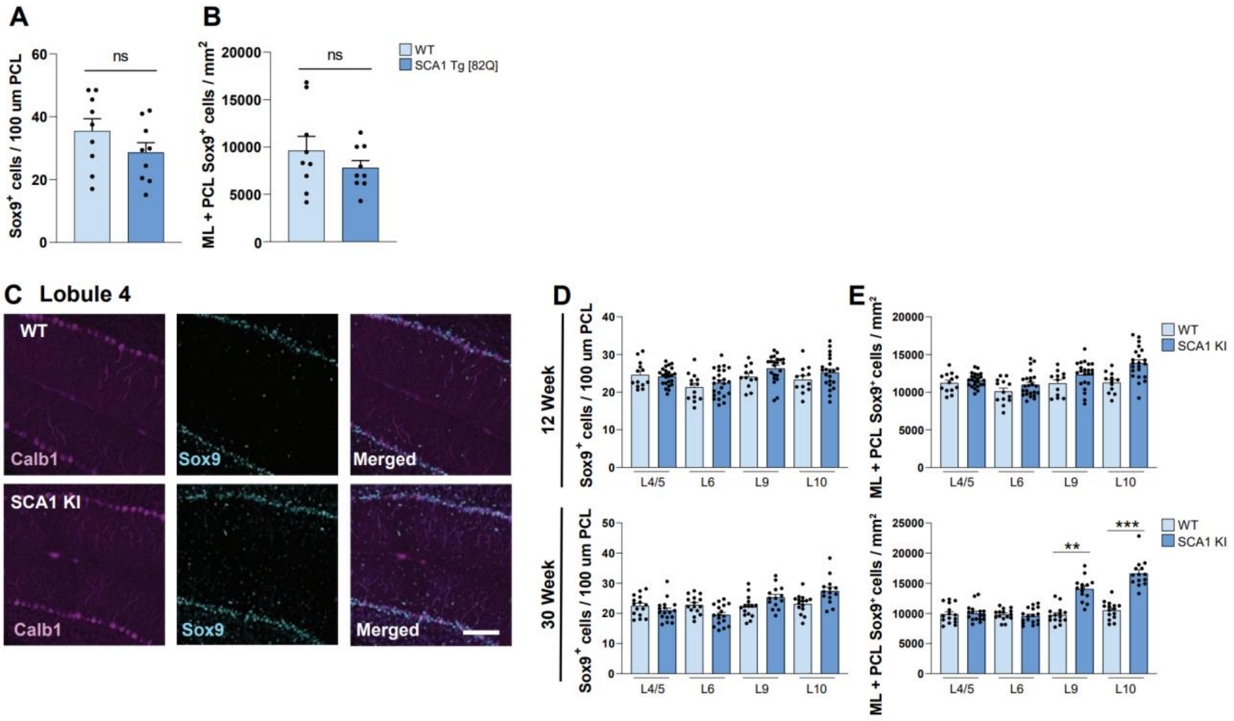


Luttik et al., Figure S4

**Figure S4,** ***Supplementary to Figure 6;*** **BG mislocalization phenotypes in SCA1 mouse models does not significantly impact BG number in PCL or total BG number.**
**A,B,** Quantification of $Sox9^+$ BG in PCL **(A)** and total in ML + PCL **(B)** in 20 week SCA1 Tg [82Q] and WT controls, cerebellar lobule L4/5**. C,** Representative images of immunostaining for $Calb1^+$ PCs and $Sox9^+$ BG in 30-week WT and SCA1 KI cerebellar lobule 4/5. Scale bar 100µm. **D,E,** Quantification of $Sox9^+$ BG in PCL **(D)** and total in ML + PCL **(E)** in 12-week (top row) and 30-week (bottom row) cerebellar lobules L4/5, L6, L9, and L10 for SCA1 KI and WT controls. $^{**}P<0.01$, $^{***}P<0.001$, ns, non-significant, by student's *t*-test (**A**,**B**), and by one-way ANOVA with Tukey's post-hoc analysis (**D**,**E**).
